# Pooled amplicon sequencing for characterizing mutations in the praziquantel molecular target TRPM_PZQ_ in schistosome populations from Western Kenya

**DOI:** 10.64898/2026.08.04.742841

**Authors:** Peter Olilah, Frédéric D. Chevalier, John Oguso, Erick Oyugi, Benjamin H. Opot, Madison Morales, Winka Le Clec’h, Tim Anderson, Eric M. Ndombi

**Author notes:** Corresponding authors: Timothy JC. Anderson, Eric M. Ndombi.

## Abstract

Mass drug administration (MDA) using Praziquantel is central to efforts to eliminate Schistosomiasis. However, regions which respond poorly to MDA (“persistent hotspots”) have been reported in many regions of Africa, including in Western Kenya. One possible explanation for persistent hotspots is that these areas contain PZQ resistant schistosome parasites. Recent studies have shown that *Sm.TRPM_PZQ_* gene is the molecular target for PZQ in schistosome parasites and that mutations in this gene can result in PZQ resistance. This study characterized mutations within *Sm.TRPM_PZQ_* in 23,420 miracidia collected from both hotspot and non-hotspot villages in Siaya County, western Kenya. We collected triplicate pools of 780.67 (SD ± 183.47) miracidia from 135 people in five hotspot villages, where *S. mansoni* prevalence remains high despite over 5 annual treatments, and from 62 people from 5 non-hotspots villages where annual treatment resulted in reduction in prevalence. We extracted DNA from each miracidia pool, amplified 15 amplicons covering 1,695bp of the *Sm.TRPM_PZQ_*transmembrane domain and sequenced these to high read depth (21,110x) using a Miseq at KEMRI-CGHR. We identified five high confidence (frequency ≥ 0.01) *Sm.TRPM_PZQ_* variants. These included four synonymous changes and a non-synonymous variant (p.L1476I). p.L1476I is found at similar frequency in non-hotspot (0.040 ± 0.006) and hotspot villages (0.044 ± 0.0050) (Mann Whitney U=18, p= 0.31) and does not impact PZQ-response in Ca^2+^ reporter assays. Our studies show that resistance variants in *Sm.TRPM_PZQ_* are rare or non-existent in the locations studied and do not explain the existence of hotspots in this region.

## INTRODUCTION

Mass treatment with praziquantel monotherapy is the foundation of schistosomiasis control. Currently, ∼250 million treatment doses are administered each year (Deol et al., 2019). This has enormous public health benefits reducing infection levels and transmission, and importantly, the proportion of people with heavy infections and severe schistosomiasis pathology. However, praziquantel treatment is imperfect: cure rates, assessed by egg counts are 73.6% (95% CI: 63.5–81.40) for *S. haematobium* and 76.4% (95% CI: 71.5–81.0) *S. mansoni*, while egg reduction ratios (ERR) are 94.7% (95% CI: 92.7–96.4) for *S. haematobium* and 95.3% (95% CI: 94.2–96.2) for *S. mansoni* (Zwang & Olliaro, 2017). Praziquantel does not kill immature parasites (Colley et al., 2014), hence maturation of immature worms after praziquantel treatment many partially explain the imperfect cure rates observed. Resistance of adult worms to praziquantel treatment is an alternative and worrying possibility.

Schistosomiasis hotspots, where high levels of infection are maintained despite annual chemotherapy, have been observed in several African locations, including Côte d’Ivoire, Kenya, Mozambique, and Tanzania (Kittur et al., 2019, 2020; Lim et al., 2023) and are a barrier to effective control. One well studied hotspot is in Western Kenya (Wiegand et al., 2017) where hotspot villages maintained higher levels of infection than non-hotspot villages even after 8 years of treatment (Olilah et al., 2026). The biological explanations for the existence of hotspots are currently unknown. One possible explanation is that these locations have extremely high transmission levels and so rapidly regain pretreatment infection levels. In Kenya, particular snail species are associated with hotspots consistent with environmental determinants (Mutuku et al., 2019). A second possibility is that such hotspots represent regions of emerging praziquantel resistance.

Recent advances in our understanding of the mechanism of action of praziquantel now make molecular screening for resistance possible (Marchant, 2024). Praziquantel binds with a transient receptor potential channel (TRPM_PZQ_) encoded by a gene on chromosome 3 of the schistosome genome: channel activation results in influx of rapid Ca^2+^ influx into cells, muscle contraction, and tegument damage, leading to worm death. Point mutations in specific sites within the S1, S2, S3, S4 helices and the TRP box – a 1,695 bp region comprising gene exons 22, 23, 27, and 31 in the isoform 5 TRPM_PZQ_ transcript – can prevent channel activation(Park et al., 2021). One of these point mutations is found naturally in a related trematode (liver fluke, *Fasciola hepatica*) (Rohr et al., 2023) making these parasites naturally resistant to PZQ. Hence, screening for point mutations in the TRPM_PZQ_ S helices and TRP-box provides a promising approach to screen schistosome populations for mutations underlying emerging praziquantel resistance. In addition, there is now a growing database of the mutations that prevent activation of the TRPM_PZQ_ when this channel is expressed in HK293 cells (https://www.trptracker.live/) (Rohr et al., 2026), which allows direct evaluation of the impact of mutations identified in the field.

One screening option involves sequencing of the TRPM_PZQ_ from individual miracidia collected from urine or fecal samples. Two studies have used genome or exome sequencing data from single miracidia for this purpose. Le Clec’h et al. (Le Clec’h et al., 2021) examined 259 miracidia from 5 countries and found one parasite with heterozygous mutations predicted to result in a truncated TRPM_PZQ_ protein and lead to channel inactivation. Berger et al. (Berger et al., 2026) examined genome sequence data from 570 miracidia and identified 4 putative praziquantel resistance mutations, which eliminated or reduced channel activation in *in vitro* Ca^2+^ reporter assays. The two mutations (p.Y1554C and p.Q1670K) that eliminated TRPM_PZQ_ activation in Ca^2+^ reporter assays were found in heterozygous state in single miracidia. False positive base calls are a particular concern when SNPs are seen in only one individual, which is the case for all putative resistance SNPs identified to date (see also (Le Clec’h et al., 2021). Such singleton calls are typically removed from population genetics studies as they are deemed “low confidence”. Hence, to date, there is no unambiguous evidence for PZQ-resistance alleles in nature.

Pooling samples provides an efficient and cost-effective approach for scaling up surveillance efforts to examine mutations within TRPM_PZQ_. Pooling has been used extensively in free living organisms for accurate measurement of allele frequencies in genome scale studies (Schlötterer et al., 2014). Likewise, pooling has seen increasing use in parasite genomics and diagnostics. For example, pooled genome sequencing allows accurate measurement of allele frequencies for bulk segregant genetic mapping in malaria parasites (Cheeseman et al., 2015; X. Li et al., 2022), and in helminth parasites (Chevalier et al., 2014; Doyle et al., 2019, 2022). The design of this project has particularly strong parallels with amplicon deep sequencing approaches used for large scale evaluation of SNPs underlying anthelminthic resistance in nematode populations: Avramenko et al. (Avramenko et al., 2019) used deep sequencing of the β-tubulin gene of nematode larvae from 194 sheep herds to examine allele frequencies of 3 SNPs underlying anthelminthic resistance in 7 different helminth species.

The central aims of this project were: (i) to design amplicons across the TRPM_PZQ_ S helices and TRP-box (ii) to validate pooled sequencing approaches for allele frequency determination in *S. mansoni* and (iii) to determine frequencies of TRPM_PZQ_ mutations underlying resistance to PZQ in *S. mansoni* from Western Kenya using high depth sequencing of miracidia pools, and (iv) investigate whether putative PZQ resistance TRPM_PZQ_ SNPs show allele frequency differences in hotspot and non-hotspot villages.

## RESULTS

### Collection of pooled miracidia from hotspot and non-hotspot villages

We screened 500 people (50 from each village) and identified infected individuals from hotspot and non-hotspot villages previously identified in the SCORE study (Wiegand et al., 2017). The results from this survey reveal that hotspot villages still showed consistently higher prevalence and intensity than non-hotspot villages despite ongoing mass chemotherapy eight years after the initial description of hotspot villages in this region (Olilah et al., 2026). There were 62 infected patients from non-hotspot and 135 infected patients from hotspot villages (Table 1). A multivariate regression analysis conducted on the number of infected individuals between hotspots and non-hotspots indicated a statistically significant difference, (*adj*. *P* □ *0.001*, *adj.* PR 95%CI = 2.08 (1.65-2.66)).

**TABLE 1.**
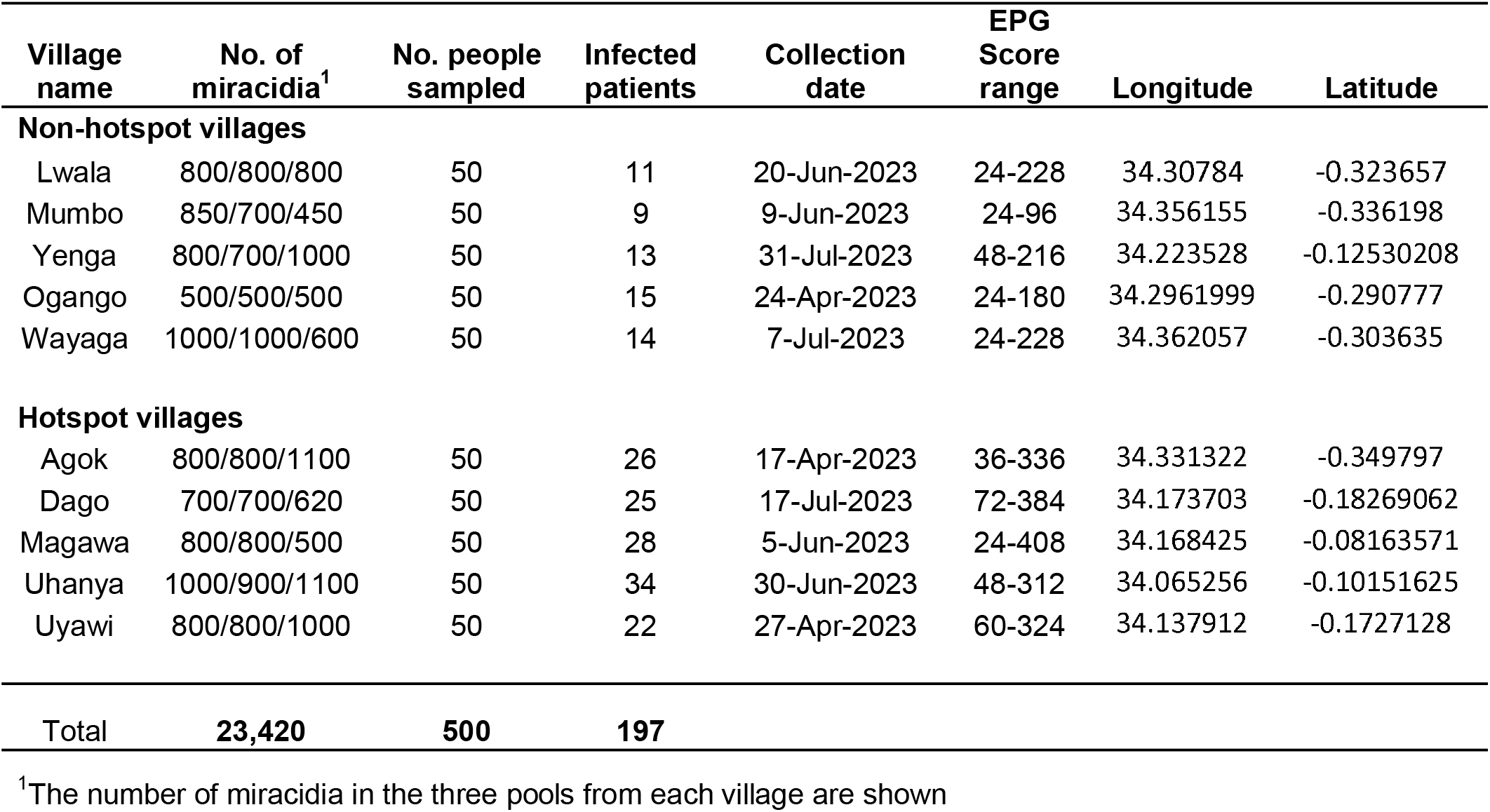
Collection of fecal samples from infected patients from non-hotspot and hotspot villages in Western Kenya.

We pooled fecal samples from infected people in each village, isolated eggs, and exposed these to sunlight in petri dishes to hatch miracidia larvae. We collected triplicate miracidia pools from each village (Table 1). The average size of each pool was 781 miracidia (range: 450-1100). In total, we collected 23,420 miracidia from the 10 villages (11,000 from non-hotspot and 12,420 from hotspot villages)

### Accuracy of pooled sequencing for allele frequency measurement

We sequenced DNA prepared from pools of 1,000 miracidia constructed from two different parasite populations (SmLE-PZQ-ER and SmLE-PZQ-ES) that differ at a single SNP (741987 T>C) within TRPM_PZQ_. These pools vary in SNP frequency from 0-10%, so can be used to quantify precision with which allele frequency can be measured using pooled sequencing. This detection can be affected by technical artifacts like index hopping (Illumina, 2017). Index hopping is characterized by exchange of indexes between reads of different libraries during the sequencing run of multiplexed libraries, with the consequence of assigning reads to the wrong library (i.e., sample) during demultiplexing. These are very rare events with minimal consequences for most applications. However, index hopping may have a significant impact in detecting and accurately estimating the frequency of rare mutations, generating false positive detection.

To evaluate optimal barcoding approaches, we ran these experiments using (i) combinatorial dual indexes (CDIs) and (ii) unique dual indexes (UDIs). We sequenced the PCR products to high read depth (140,824-540,133) on an Illumina iSeq 100 and measured the frequency of this SNP in each pool. We carried out detection of ER alleles in the controls because of index hopping. Using CDIs, we could not obtain accurate allele frequency below 0.01: our negative control (0.0063 - 0.0068) and 0.001 sample (0.0070 - 0.0081) showed observed frequency just under 0.01. However, using UDIs we obtained accurate allele frequencies down to 0.001, although we also measured low levels (0.00042 - 0.00046) of false positive mutations in the negative controls (Figure 3). We therefore opted to use UDI for examination of field samples. We also opted to use a conservative allele frequency threshold of 0.01 for detection of high confidence mutations and to exclude false positive mutations.

**Fig. 1.**
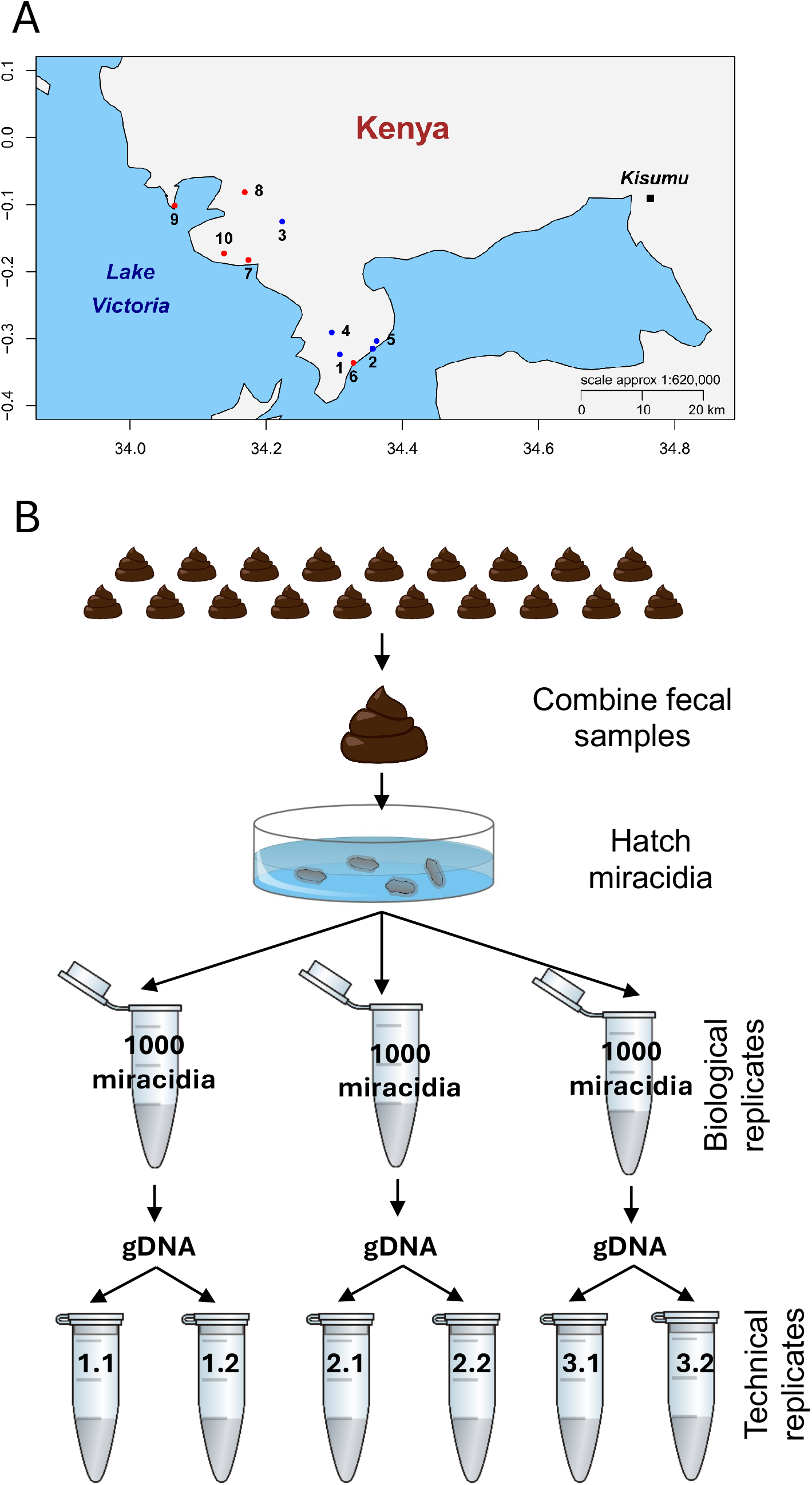
Sampling of miracidia pools. **A. Sampling locations.** We identified infected patients in 10 villages in Western Kenya. There were 5 non-hotspot villages (blue dots) (1. Lwala; 2. Mumbo; 3. Yenga; 4: Ogango; 5 Wayaga) and 5 hotpot locations (red dots) (6. Agok; 7. Dago; 8. Magawa; 9. Uhanya; 10. Uyawi). Numbers of infected individuals and GPS locations are shown in Table 1. **B. Sampling methodology.** We combined fecal samples from multiple infected people from each village. Following isolation of eggs, we hatched miracidia. Hatched miracidia were collected into three tubes containing pools of 1000 miracidia (biological replicates) from which we prepared genomic DNA. We performed PCR in parallel on two technical replicates from each biological replicate.

**Fig. 2.**
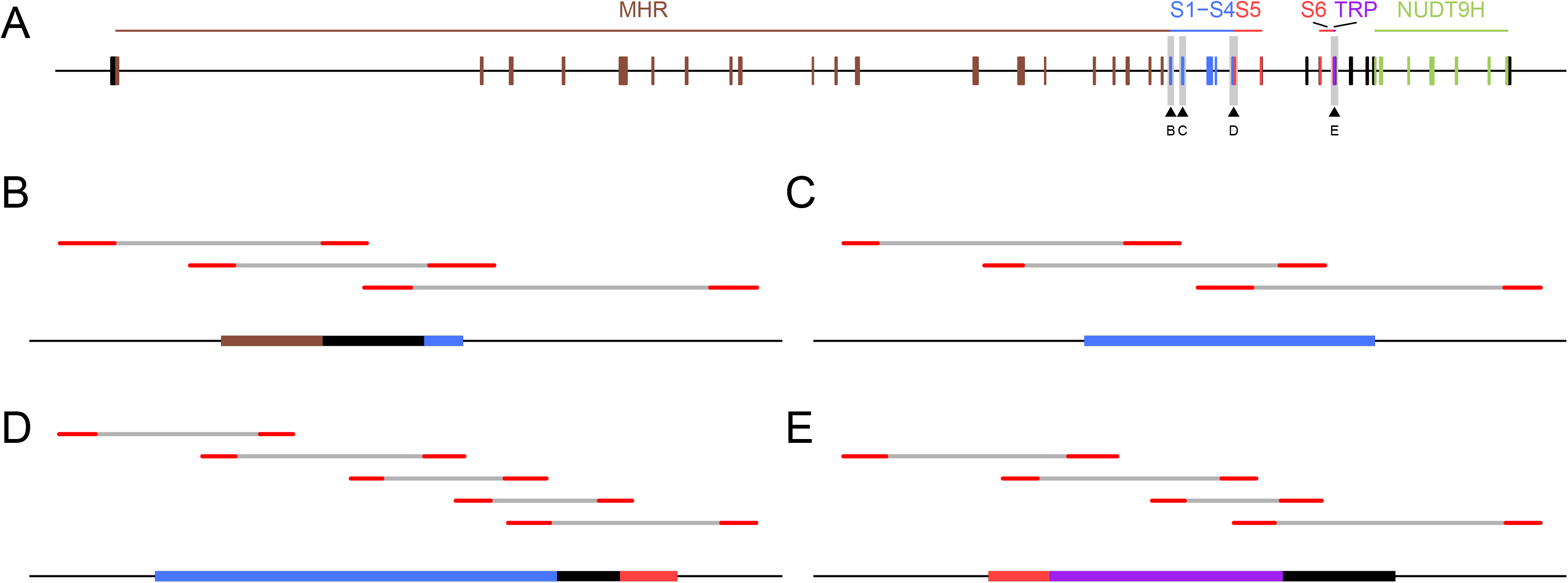
Regions of SmTRPM_PZQ_ channel gene targeted for amplicon sequencing. **(A)** Gene structure showing the position of the exons (rectangle) and the different domains they encode. The arrow heads point to the targeted exons (gene exon 22, 23, 27, 31). **(B to E)** Position of the amplicons for each exon. The red lines represent the primers, the grey line the amplified section. The thin and thick dark lines represent the intron and exon, respectively. The rectangle represents the protein domains encoded following the same color code as for (A).

**Fig. 3.**
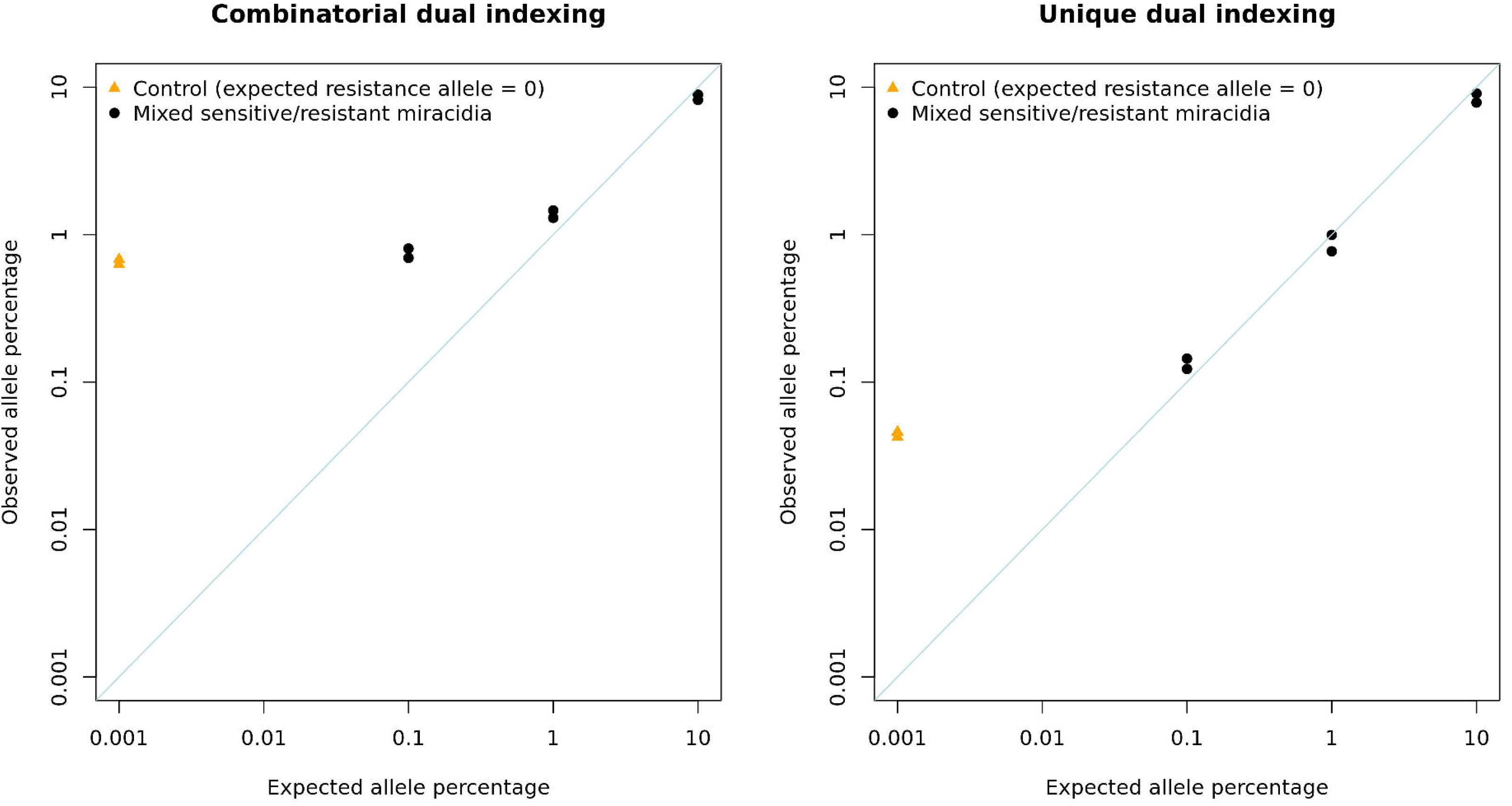
Allele frequency of known mutation using amplicon sequencing of miracidia pools. We used SmLE-PZQ-ES and ER, which differ at a single SNP (741987 T>C) within the *Sm.TRPM_PZQ_*, to generate pools at different allele frequencies (0, 0.001, 0.01, 0.1) by using different ratio of ER:ES individuals (0:1000, 1:999, 10:990, 100:900, respectively). Each pool was replicated. We amplified the locus of the SNP and performed amplicon sequencing using either combinatorial or unique dual indexes (CDI and UDI, respectively). We compared the observed allele frequency of the SNP associated to ER from each pool (dot or triangle) to the expected allele frequency. CDI showed higher than expected ER allele frequency (∼0.01) for 0 and 0.001 allele frequency, likely due to index hoping events. UDI showed superior sensitivity as it allows to accurately detect allele at 0.001 frequency, while few ER alleles were still detectable at very low frequency in the control pool (0), likely due to minor index hoping events. This demonstrates that UDI are optimal for accurate allele frequency detection from pools.

### Sequencing TRPM_PZQ_ from pooled miracidia from hotspot and non-hotspot villages

We generated 15 PCR products from the triplicate miracidia samples (biological replicates) from each village. This was done in duplicate (technical replicate) for each biological replicate. We prepared sequencing libraries from each biological and technical replicate, using UDI to index each library. The mean read depth was 11,879.89 (range: 45.99-30,600.85) across the regions sequenced (Figure 4).

**Fig. 4.**
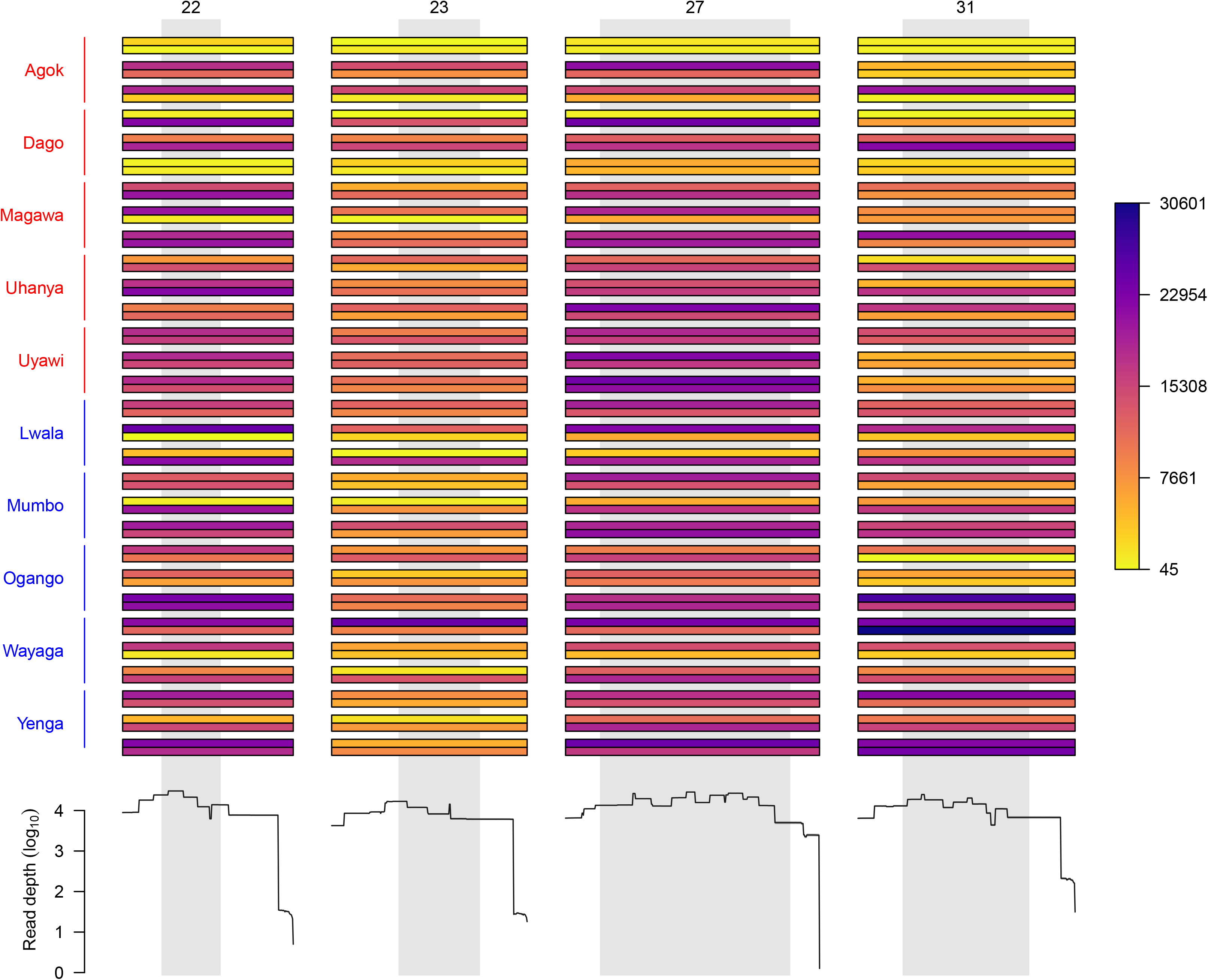
Read depth across amplicons and samples from pools of field miracidia. Collated bars represent technical replicate while group of three collated bars represent the biological replicate for each village (red = hotspot, blue = Non-hotspot). The color of the bar represents the read depth as indicated in the legend. The bottom section represents the average read depth at each position across all samples. The grey section highlights the exon sequence. Overall, a large proportion of the samples shows high read depth, especially on the exon region, which allows for the accurate detection of low frequency mutation.

We filtered sequence data to remove sites (i) with read depth <100; (ii) alternative allele read depth <2; (iii) extreme strand bias (SOR<2); (iv) high variance in allele frequency in technical replicates; (i) average allele frequency <0.01 (see methods for details). We used these filtering criteria to identify high confidence SNPs; both the scripts and the sequence data are available for those who want to investigate the dataset using alternative filtering approaches.

After filtering we found 5 high confidence variants: one non-synonymous variant (g.118263T>A, p.L1476I) and four synonymous variants (g.118178C>T, g.128912C>T, g.128924T>C and g.128969A>G). Four of the SNPs were at comparable frequency in the 10 villages: p.L1476I (freq = 0.043 ± 0.005 (S.D.)), g.128912C>T (0.049 ± 0.010), g.128924T>C (0.051 ± 0.008) and g.128969A>G (0.051 ± 0.023). One of the synonymous SNPs (g.118178C>T) was only found in >1% of reads in one pooled sample from one village. Figure 5 shows the allele frequencies of these five SNPs in non-hotspot and hotspot villages. Allele frequencies were similar across all villages for p.L1476I and g.128924T>C. However, g.128969A>G showed high variance, which resulted from very low frequencies in one non-hotspot village (Lwala), and g.118178C>T showed very low frequencies except in one village where it reached 0.01 (Yenga) T-tests revealed no difference in allele frequencies between non-hotspot and hotspot villages for any of the five SNPs (g.118178C>T: Mann Whitney U=6, p=0.22; p.L1476I: U=18, p=0.31; g.128912C>T: U=11, p=0.84; g.128924T>C: U=12, p=1.0; g.128969A>G, U=7, p=0.31).

**Fig. 5.**
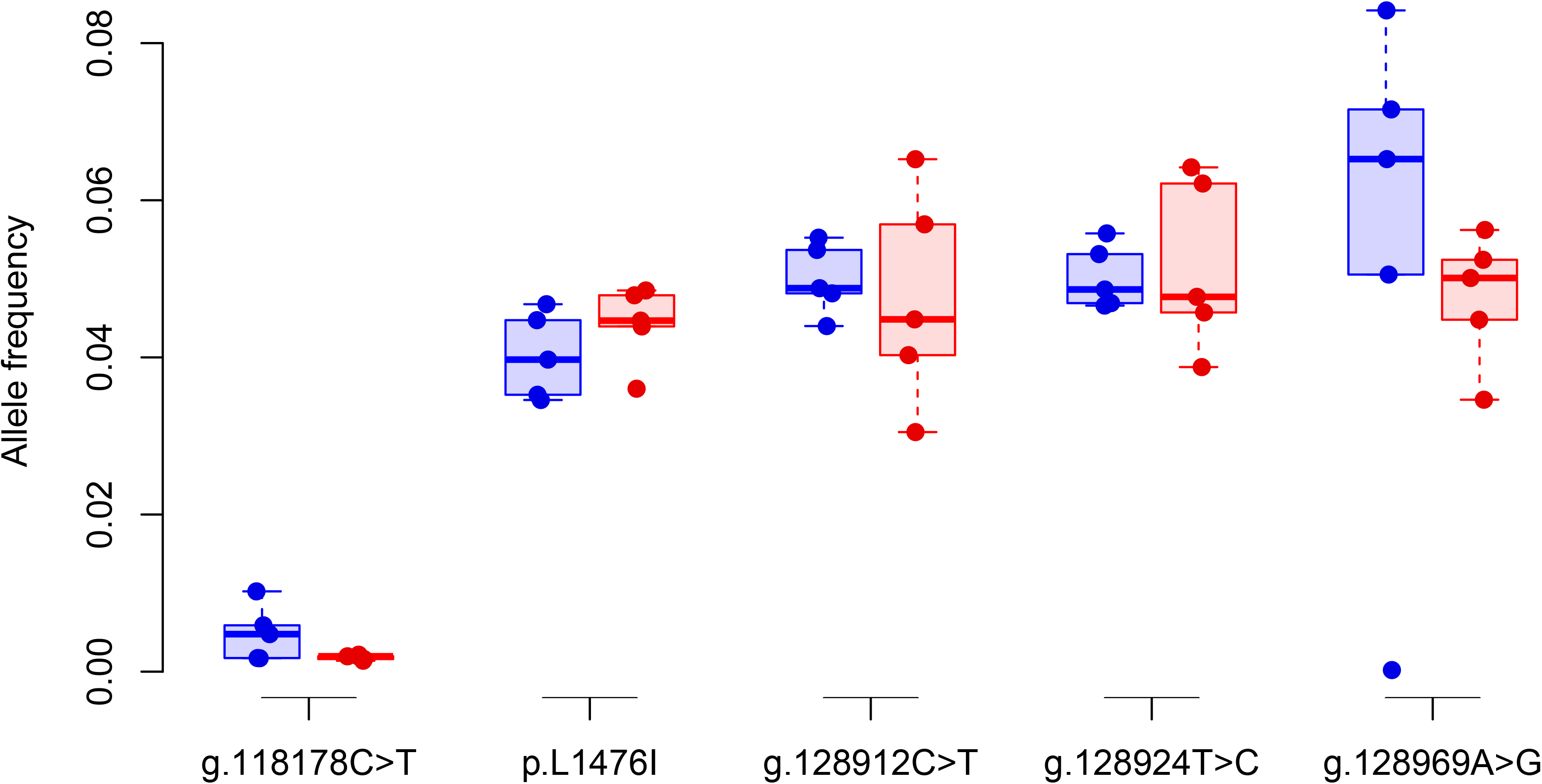
Allele frequency of mutations identified from pools of field miracidia. We identified five mutations, one non-synonymous (g.118263T>A, p.L1476I) and four synonymous (g.118178C>T, g.128912C>T, g.128924T>C, g.128969A>G). The first mutation has an average frequency of 0.003 with an outlier at 0.01. The other four mutations have an allele frequency ranging from 0.03 to 0.08 with an outlier at 0.002 Each mutation has a similar frequency between Non-hotspot (blue) and hotspot (red). Dots represent the allele frequency of a mutation for a location.

## DISCUSSION

We used deep sequencing of amplicons in pools of miracidia to screen 23,420 parasite genomes for variants-of-concern within the TRP-Box region of TRPM_PZQ_ gene. We observed just five high confidence variants (p.L1476I and three synonymous mutations), that were scored in technical replicates from miracidia pools and had allele frequency ≥ 0.01 in at least one miracidia pool. p.L1476I was present in all 10 populations examined with mean frequency of 0.043 (± 0.005 (S.D.)) and we observed no difference in the allele frequency in hotspot and non-hotspot villages. The p.L1476I SNP was previously observed at low frequency (frequency = 0.04) in Tanzania (Le Clec’h et al., 2021) as wells as in Uganda (Berger et al., 2026). This mutation is situated within the extracellular loops (S3 and S4) of the channel and does not impact PZQ response *in vitro* measured with the Ca^2+^ reporter assay (TRPtracker: https://www.trptracker.live/) (Berger et al., 2026). Three synonymous mutations were also present at moderate frequency (g.128912C>T: 0.049 ± 0.010, g.128924T>C: 0.051 ± 0.008; g.128969A>G: 0.051 ± 0.023). These three mutations were previously identified in an exome sequence survey from single miracidia: g.128912C>T (frequency = 0.045) and g.128924T>C (frequency = 0.031) were found in Tanzania, while g.128969A>G was found in Tanzania (frequency = 0.11) and the middle East (frequency = 0.050) (Le Clec’h et al., 2021).

The five mutations we observed in the TRPM_PZQ_ in Western Kenya are not variants of concern. These molecular data suggest that mutations in the transmembrane spanning regions and the TRP-box of TPRM_PZQ_ are unlikely to result in PZQ-R in *S. mansoni* from Western Kenya. Furthermore, the absence of difference in allele frequency in hotspot and non-hotspot villages is consistent with the conclusion that other factors may drive the consistent differences in transmission in the face of repeated mass chemotherapy with PZQ.

Our study targeted the PZQ binding region of TRPM_PZQ_ containing the transmembrane spanning regions and the TRP-box of TPRM_PZQ_ because this region interacts with PZQ, and *in vitro* tests of engineered mutations show that many amino acid changes in this gene region can impact response to PZQ (Rohr et al., 2026). However, the TRPM_PZQ_ gene is large, and it is possible that mutations in other regions of this gene may impact PZQ-response. Consistent with this, two mutations (F107A and G108A) engineered in the NH_2_-terminus of TRPM_PZQ_ prevent channel activation by PZQ *in vitro* (Rohr et al., 2026). Further examination of engineered mutations and natural mutations outside the TRP-box may result in the discovery of further mutations outside the gene regions examined that can impact resistance.

While we cannot therefore eliminate the possibility that mutations outside the PZQ binding region impact PZQ-R, we believe that this is unlikely. In a parallel study conducted in 6 of the 10 villages described here, we examined *in vitro* PZQ-sensitivity of adult worms derived from each village (Ndombi et al., 2026). This was done by hatching eggs, infecting snails with miracidia, and infecting hamsters to generate large numbers of schistosome genotypes from each village. We then directly examined PZQ-response in these field derived parasites using a movement assay. These lab intensive studies indicated that PZQ-R is rare or non-existent in *S. mansoni* from Western Kenya, consistent with the conclusions from our molecular study.

### Limitations

The molecular methods we used involved individual amplification of 15 PCR products from each sample to cover the region of interest, and quantification to generate equimolar mixtures for Illumina sequencing. This was done to ensure even sequence coverage across the gene regions targeted. We suggest that future approaches should optimize single tube amplicon panels to simplify library preparation (Campbell et al., 2015). Single tube amplicon panels could also be expanded to include the complete TRPM_PZQ_ sequence as well as other loci of interest, such as the sulfotransferase (SULT-OR) locus underlying oxamniquine resistance (Chevalier et al., 2019; Valentim et al., 2013). An alternative approach would be to use RNA baits and targeted enrichment (Chevalier et al., 2014); these methods typically allow large numbers of loci to be screened than amplicon panels.

We used a minimum frequency threshold of 0.01 for scoring mutations in TRPM_PZQ_. This is equivalent to detecting 20 mutant alleles in a pool of 1000 miracidia (or 2000 genomes). Hence, our pooling approach minimizes scoring of false positive mutations but may not detect very rare alleles.

## MATERIALS AND METHODS

### Ethics

The laboratory study was performed in accordance with the Guide for the Care and Use of Laboratory Animals of the National Institutes of Health. The protocol was approved by the Institutional Animal Care and Use Committee of Texas Biomedical Research Institute (permit number:1419-MA).

The field study was approved by the Kenya Medical Research Institute Scientific and Ethical Review Unit (SERU Protocol no. 3218) and the Kenyatta University Ethics Review Committee (Protocol no. PKU/2206/I1352). Participants were asked to provide informed consent before enrollment in the study, while for minors, parental informed consent was obtained.

### Field study design and sample collection

This was a cross-sectional study carried out in Siaya County, western Kenya in five ‘persistent hotspot’ and five ‘non-hotspot villages’ of schistosomiasis transmission near Lake Victoria. Persistent hotspots are villages whose *S. mansoni* prevalence did not significantly reduce following praziquantel mass drug administration (Wiegand et al., 2017). These 10 are among villages identified previously (Wiegand et al., 2017). A total of 500 participants were recruited from five persistent hotspots and five non-hotspots. For diagnosis of schistosome infection, one fecal sample was collected from each participant for the Kato-Katz test (Altman et al., 2025), using two slides per sample for determining the number of eggs per gram of stool. The participants who were *S. mansoni* positive from the fecal sample collected were recruited for the collection of larger stool samples in 100 ml containers. The collection and processing of the fecal samples was done by an experienced technician in the parasitology laboratory in the Neglected Tropical Diseases Unit at the Kenya Medical Research Institute’s Centre for Global Health Research (KEMRI-CGHR). Following diagnosis, the additional large fecal samples collected from each recruited participant were used for miracidia hatching to prepare pools of miracidia. Each positive participant received a dose of praziquantel.

To obtain schistosome larvae, approximately 100g of fecal sample from each individual patient of a village were mixed and processed through 3 layers of sieves (250μm, 90μm, and 45μm). Isolated eggs were then placed in petri dishes containing freshwater and exposed to light to allow miracidia hatching. We collected triplicate pools of miracidia (450-1100 per pool) in 1.5 mL tubes from each of five hotspot and five non-hotspot villages. Each pool was kept at - 20°C until DNA extraction.

### Laboratory study design to test combinatorial and unique dual index

We used SmLE-PZQ-ES and SmLE-PZQ-ER, which differ at a single SNP (741987 T>C) within the *Sm.TRPM_PZQ_*, to generate pools at different allele frequencies (0, 0.001, 0.01, 0.1) by using different ratios of ER:ES individuals (0:1000, 1:999, 10:990, 100:900, respectively). Each pool was replicated. We collected miracidia from eggs recovered from the livers of infected hamsters. Briefly, we minced the livers, blended them for 1 minute, filtered the homogenate using a gauze to remove large particles then using a 100 µm nylon mesh to retain the eggs. The filter was washed with freshwater to collect the eggs and the solution was placed under light for 1h. Miracidia were collected in ∼ 1mL under a microscope using an elongated glass pipette to minimize the volume sampled. Pools were stored at -20°C until DNA extraction.

### DNA extraction of miracidia pools

We extracted gDNA from miracidia pools using the DNeasy Blood & Tissue kit (Qiagen) following the manufacturer’s protocol. Briefly, we thawed the miracidial pool and centrifuged it at 800 × *g* for 5 minutes to pellet the miracidia and remove most of the water. We lysed the miracidia for 1h at 56°C as per the protocol, passed the sample on the column, washed it twice before eluting the DNA in 50 µL (laboratory pools) or 200 µL (field pools) of ultrapure water. We quantified the extracted gDNA using the Qubit dsDNA High Sensitivity Assay kit (Thermo Fisher Scientific). Samples were stored at -20°C until use.

### Two-step amplicon libraries

#### a) First step: PCR amplification of targeted exons

We added partial adapter sequences on the 5’ end of all primer used for amplification to allow the attached of the indexes for the second step. We used Nextera partial adapter sequences for combinatorial dual indexes (CDIs) (forward direction: 5’–TCGTCGGCAGCGTCAGATGTGTATAAGAGACAG–3’, reverse direction: 5’–GTCTCGTGGGCTCGGAGATGTGTATAAGAGACAG–3’) and TruSeq partial adapter sequences for unique dual indexes (UDIs) (forward direction: 5’–ACACTCTTTCCCTACACGACGCTCTTCCGATCT–3’, reverse direction: 5’–GTGACTGGAGTTCAGACGTGTGCTCTTCCGATCT–3’).

For laboratory test pools, we amplified a single locus carrying the mutation of interest using KAPA HiFi HotStart ReadyMix PCR kit (Roche). For each pool, we used 20 ng of DNA template, 0.75 µL of forward and reverse primers (10 µM), 12.5 µL of master mix and the volume of ddH_2_O required for a final volume of 25 µL. The locus-specific primer sequences are 5’–CAACAACACCAATTCAATCAC–3’ (forward) and 5’–AAGTACTGGTTTATTTGATTTCTC–3’ (reverse). We used the following amplification program: 95°C for 3 minutes, [95°C for 30 seconds, 55°C for 30 seconds, 72°C for 30 seconds] × 25, 72°C for 5 minutes. We visualized PCR products on 2% agarose gel.

For miracidia pools from each village, we carried out amplicon PCR preparation in 2 technical replicates per miracidia pool using 15 Sm.TRPM_PZQ_ primers and High Fidelity PCR EcoDry premix (Takara) following the manufacturer’s protocol. Each PCR mix included 21 µL of rehydrated master mix with ddH_2_O, 0.5 µL of each forward and reverse primers (10 µM), and 3 µL of DNA sample. We used the following amplification program: 95°C for 3 minutes, [95°C for 30 seconds, primer T_m_ for 30 seconds, 72°C for 30 seconds] × 35, 72°C for 5 minutes. Primer sequences and associated T_m_ are available in Table 2, full primer sequences are available in Supp. Table 2. We visualized PCR products on 2% agarose gel.

**TABLE 2:**
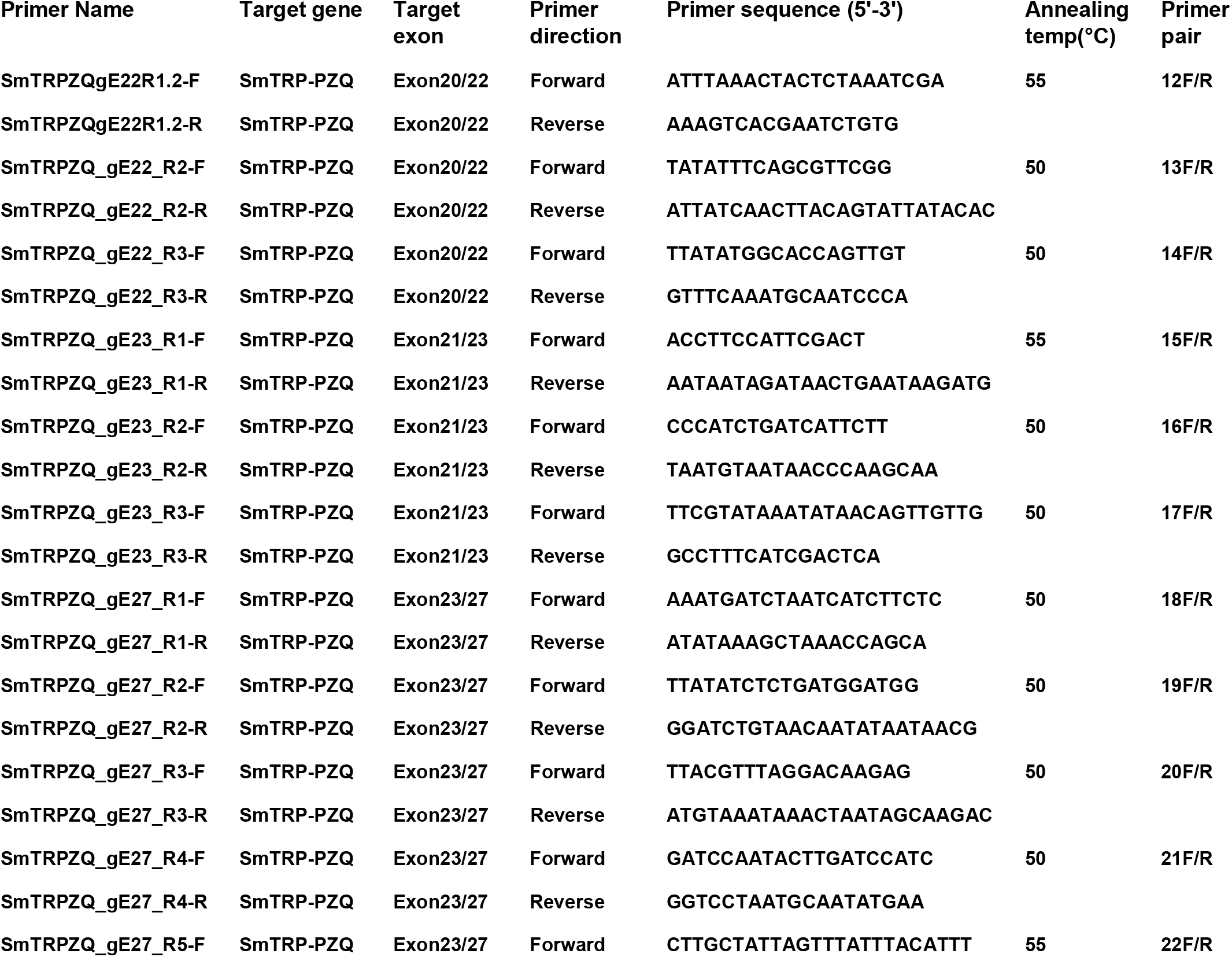
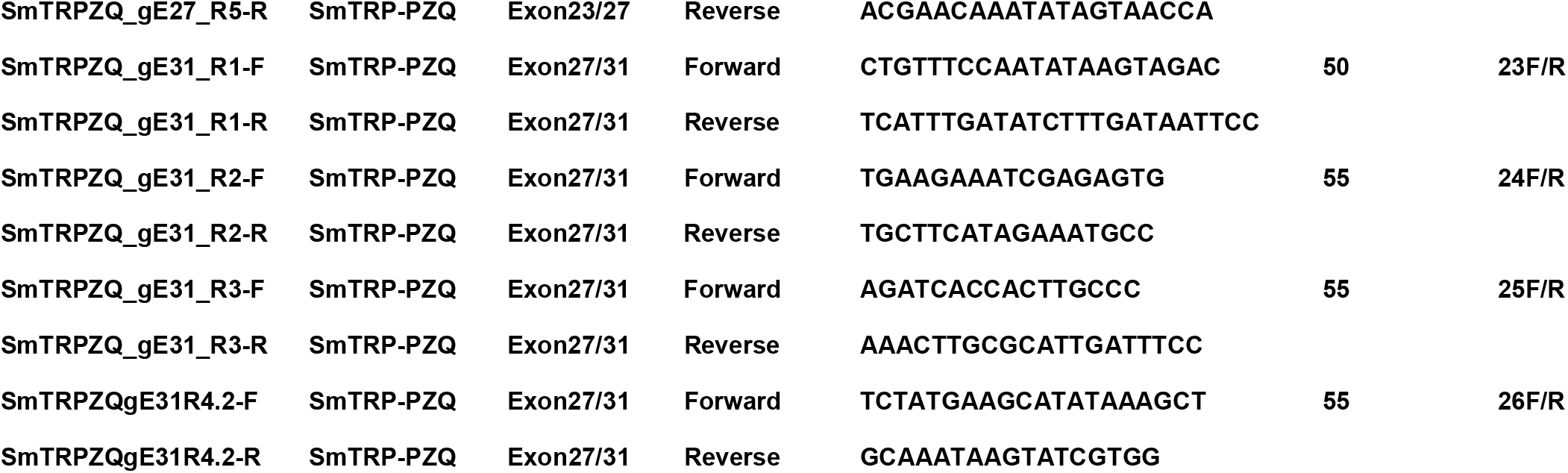
Genomic locations, sequence, and annealing temperature of primers used to amplified exon 22, 23, 27 and 31 of the *Sm.TRPM_PZQ_*. We added TruSeq partial adapters sequences on their 5’ end of all primers. See Materials and Methods for details.

All PCR products were purified using KAPA pure beads (Kapa Biosystems, South Africa) with beads to sample ratio of 3X following the manufacturer’s protocol. We eluted the cleaned PCR amplicons using 52.5 µL of PCR-grade water to recover 50 µL. We then quantified each of the laboratory PCR products using Qubit dsDNA High Sensitivity Assay kit. The field PCR products were quantified using Invitrogen Picogreen assay kit (Thermo Fisher Scientific) and Tecan fluorescent reader (Tecan).

#### **b)** Second step: Library indexing

We added indices to each 5’ end of PCR amplicons to trace the sequencing data to each sample. The laboratory amplicon libraries were indexed either using combinatorial dual indexes (CDIs) or unique dual indexes (UDIs). The same DNA template was used for the testing of the two methods. We ordered custom Nextera CDI primers from Eurofins and pre-made UDI primers from Integrated DNA Technologies (IDT, xGen™ UDI Primer Plate 1, 8nt). We indexed each library by PCR using KAPA HiFi HotStart ReadyMix. We used 5 µL of DNA template, either 2.5 µL of P5 and P7 CDI primers (10 µM) or 5 µL of UDI primers, 25 µL of master mix and 15 of ddH2O for a final volume of 50 µL.

Each field library was generated by pooling 100 ng of each of the 15 PCR products of a given sample. With 3 biological replicates per village, and 2 technical replicates per biological replicate, we had a total of 60 libraries. We performed PCR to index each library using High Fidelity PCR EcoDry premix (Takara) and UDIs (xGen™ UDI Primer Plate 1, 10nt, IDT) (Wisely, 2025). Each PCR mix included 25 µL of rehydrated master mix with ddH_2_O, 10 µL of ddH_2_O, 10 µL of library pool, and 5 µL of UDI for a final volume of 50 µL.

For all CDI and UDI libraries, we used the following amplification program: 95°C for 3 minutes, [95°C for 30 seconds, 55°C for 30 seconds, 72°C for 30 seconds] × 8, 72°C for 5 minutes. We then performed a PCR clean-up on each indexed library using KAPA beads as described in the previous section, using a 1.5X beads to sample ratio. We eluted the cleaned PCR amplicons using 27.5 µL of PCR-grade water to recover 25 µL. We determined the size and concentration of each library using Tapestation D1000 Screen Tape Assay (Agilent Technologies) and made an equimass sequencing pool by pooling 200 ng of each library.

#### **c)** Sequencing

For laboratory libraries, we quantified the molarity of each library using KAPA Library Quantification Kit (Roche). We made an equimolar pool of libraries for sequencing. The sequencing pool was mixed with 25% PhiX and sequenced on an Illumina iSeq100 at Texas Biomedical Institute. Raw sequencing data of laboratory libraries are accessible from the NCBI Sequence Read Archive under BioProject accession number PRJNA1405726

For the field sequencing pool, we quantified the molarity of the final pool by qPCR using QIAseq^TM^ Library Quant Assay kit (Qiagen, USA). The final pool was mixed with 20% of PhiX and then paired-end sequenced on a single flow-cell lane of an Illumina MiSeq sequencer at the Kenya Medical Research Institute’s Centre for Global Health Research (KEMRI-CGHR) in Kisumu. Raw sequencing data of field libraries are accessible from the NCBI Sequence Read Archive under BioProject accession number PRJNA1406309.

#### Variant analysis of targeted exons of *Sm.TRPM_PZQ_*

We used Jupyter notebook and scripts for processing the sequencing data and identifying variants (https://github.com/fdchevalier/PZQ_allele_suveillance_Kenya). We trimmed primer sequences from the sequencing data using Cutadapt (v3.5) (Martin, 2011). We aligned the trimmed sequencing data against the *S. mansoni* reference genome v7 using BWA (v0.7.17) (H. Li & Durbin, 2009). We indexed the aligned data using SAMtools (H. Li et al., 2009). We performed joint variant calling for indels and single nucleotide variants using FreeBayes (v.1.3.6) (Garrison & Marth, 2012). We normalized, split and atomized multi-allelic sites and complex variants using BCFtools (Danecek et al., 2021). We evaluated the read depth of each amplified locus using BEDtools (v2.30.0) (Quinlan & Hall, 2010). We evaluated the functional impact of each variant using a custom script (Chevalier et al., 2019).

We performed data quality filtering and allele frequency analysis using R (v4.1.3)(R Core Team, 2021). We kept only biallelic variants and then filtered out variants showing strand odd ratio (SOR) greater than 2. We removed genotypes with allele frequencies lower than the minimal expectation (i.e., 1 allele / [number of miracidia × 2]), with total read depth lower than 100, and with alternative allele depth lower than 2. We filtered out sites that were no longer variable. We then filtered out genotypes based on technical replicates. We first removed genotypes with an average allele frequency in the technical replicates <0.01. We then removed genotypes showing high variation between replicates by transforming the allele frequency using arcsine-square-root and excluding genotypes showing a difference in transformed allele frequency greater than 0.1 between replicates.

### Statistics

We tested if allele frequencies of variants were different between hotspot and non-hotspot villages using a Mann-Whitney test. Results were considered statistically significant when p-values < 0.05. Statistical tests were perform using R (v4.1.3).

## Availability of data and materials

Raw sequencing data are accessible from the NCBI Sequence Read Archive under BioProject accession numbers PRJNA1406309 and PRJNA1405726. Commands and scripts used for processing sequencing data and performing downstream analysis are available in a Jupyter notebook on Github (https://github.com/fdchevalier/PZQ_allele_suveillance_Kenya). Final VCF files are available on Zenodo (DOI: 10.5281/zenodo.21776379)

## Authors’ contributions

FDC, WL, TJCA, and EN designed the experiments. MM, FC and WL validated the primers. JO, EO and PO prepared and collected the field samples, extracted DNA and prepared the amplicon libraries. BHO coordinated sequencing run in Kenya. FDC performed the data analyses. PO, FDC, TJCA and EN wrote the first version of the manuscript. All authors edited the manuscript and approved the final version.

## Acknowledgements

We thank the participants and community health workers from the 10 villages for their involvement with this project. Clint Christensen and Jeremy Glenn at the Texas Biomed Core for assistance with sequencing libraries.

